# EEG mismatch responses in a multi-modal roving stimulus paradigm provide evidence for probabilistic inference across audition, somatosensation and vision

**DOI:** 10.1101/2022.10.27.514010

**Authors:** Miro Grundei, Pia Schröder, Sam Gijsen, Felix Blankenburg

## Abstract

The human brain is constantly subjected to a multi-modal stream of probabilistic sensory inputs. EEG signatures, such as the mismatch negativity (MMN) and the P3, can give valuable insight into neuronal probabilistic inference. Although reported for different modalities, mismatch responses have largely been studied in isolation, with a strong focus on the auditory MMN. To investigate the extent to which early and late mismatch responses across modalities represent comparable signatures of uni- and cross-modal probabilistic inference in the hierarchically structured cortex, we recorded EEG from 32 participants undergoing a novel tri-modal roving stimulus paradigm. The employed sequences consisted of high and low intensity stimuli in the auditory, somatosensory and visual modalities and were governed by uni-modal transition probabilities and cross-modal conditional dependencies. We found modality specific signatures of MMN (∼100-200ms) in all three modalities, which were source localized to the respective sensory cortices and shared right lateralized pre-frontal sources. Additionally, we identified a cross-modal signature of mismatch processing in the P3a time range (∼300-350ms), for which a common network with frontal dominance was found. Across modalities, the mismatch responses showed highly comparable parametric effects of stimulus train length, which were driven by standard and deviant response modulations in opposite directions. Strikingly, the P3a responses across modalities were increased for mispredicted compared to predicted and unpredictable stimuli, suggesting sensitivity to cross-modal predictive information. Finally, model comparisons indicated that the observed single trial dynamics were best captured by Bayesian learning models tracking uni-modal stimulus transitions as well as cross-modal conditional dependencies.

## Introduction

Humans inhabit a highly structured environment governed by complex regularities. The brain is subjected to such environmental regularities by a multi-modal stream of sensory inputs ultimately constructing a perceptual representation of the world. The sensory system is thought to capitalize on statistical regularities to efficiently guide interaction with the world enabling anticipation and rapid detection of sensory changes (Gregory 1980; Bregman 1994; Friston 2005; Winkler 2009; Frost 2015; Dehaene 2015).

Neuronal responses to deviations from sensory regularities can be valuable windows into the brain’s processing of statistical properties of the environment and corresponding sensory predictions. The presentation of rare deviant sounds within a sequence of repeating standard sounds induces well known mismatch responses (MMRs) that can be recorded with electroencephalography (EEG), such as the mismatch negativity (MMN; Näätänen 1978; Näätänen 2007) and the P3 (or P300; Sutton 1965; Squires 1975; Polich 2007). The MMN is defined as a negative EEG component resulting from subtraction of standard from deviant trials between ∼100-200ms post-stimulus. Although the MMN has primarily been researched in the auditory modality, similar early mismatch components have been reported for other sensory modalities, including the visual (Pazo-Alvarez 2003; Kimura 2011; Stefanics 2014) and, to a lesser extent, the somatosensory modality (Kekoni 1997; Hu 2013; Andersen 2019). The P3 is a later positive going component in response to novelty between 200-600ms around central electrodes, which has been described for the auditory, somatosensory and visual modalities and is known for its modality independent characteristics (Escera 2000; Friedman 2001; Knight 1998; Schroeger 1996; Polich 2007).

Despite being one of the most well-studied EEG components, the neuronal generation of the MMN remains subject of ongoing debate (Näätänen 2005; May 2010; Garrido 2009b). Two prominent but opposing accounts cast the MMN as adaptation-based or memory-based, respectively. Adaptation based accounts argue that the observed differences between standard and deviant responses primarily result from neuronal attenuation leading to stimulus specific adaptation (SSA; May 2004; Jääskiläinen 2004; May 1999). In animals, SSA has been shown to result in response patterns similar to the MMN (Ulanovsky 2003; 2004) and simulation work suggests that different types of MMN-like responses can be reproduced by pure adaptation models (May 2010). However, it remains unclear if the full range of MMN characteristics can be explained by adaptation alone (Garrido 2009b; Fitzgerald 2020; Wacongne 2012). The memory-based view, on the other hand, suggests that the MMN is a marker of change detection based on sensory memory trace formation (Näätänen 1990; 1992; 2005). The memory trace stores local information on stimulus regularity and compares it to incoming sensory inputs that may signal changes in the current sensory stream.

While largely neglected by previous interpretations of the MMN, it is becoming increasingly clear that key empirical features of mismatch responses concern stimulus predictability rather than stimulus change *per se*. The MMN has been reported in response to abstract rule violations (Paavilainen 2013), unexpected stimulus repetitions (Alain 1999; Horvath 2004; Macdonald 2011) and unexpected stimulus omissions (Yabe 1997; Hughes 2001; Salisbury 2012 Wacongne 2011; Heilbron 2018). Similar characteristics have been reported for P3 MMRs (Duncan 2009; Prete 2022) and both MMN and P3 responses have been shown to increase for unexpected compared to expected deviants (Sussman 1998; 2005; Schroeger 2015). Insights concerning the predictive nature of mismatch responses have led to further development of the memory-based account of MMN generation into the model-adjustment hypothesis (Winkler 2007). This view assumes a perceptual model that is informed by previous stimulus exposure and continually predicts incoming sensory inputs. The model is updated whenever inputs diverge from current predictions, and the MMN is hypothesized to constitute a marker of such divergence.

The model-adjustment hypothesis is in line with the increasingly influential view that the brain is engaging in perceptual inference to anticipate future sensory inputs (Helmholtz 1856; Gregory 1980; Friston 2005). Related theories regard the brain as an inference engine and come with neuronal implementation schemes that accomplish probabilistic (Bayesian) inference in a neurologically plausible manner (Friston 2005; 2010; Bastos 2012). Process theories such as predictive coding assume that the brain maintains a generative model of its environment which is continuously updated by comparing incoming sensory information with model predictions on different levels of hierarchical cortical organization (Rao 1999; Winkler 2012; Friston 2005; 2010). Differential influences of SSA and change detection on the MMN are proposed to result from the same underlying process of prediction error minimization, mediated by different post-synaptic changes to (predicted) sensory inputs (Garrido 2008; 2009a; Auksztulewicz 2016). As such, the theory has the potential to unify previously opposing theories of MMN generation (Garrido 2008; 2009a; 2009b) while accounting for its key empirical features (Wacongne 2012; Heilbron 2018).

With regards to the proposed universal nature of predictive accounts of brain function, reports of comparable mismatch responses across different modalities are of particular interest. So far, mismatch signals have been primarily studied in isolation, with a strong focus on the auditory system. However, key properties of the auditory MMN, such as omission responses and modulations by predictability, have also been reported for the visual (Czigler 2006; Kok 2014) and the somatosensory MMN (Naeije 2018; Andersen 2019), and modelling studies in all three modalities suggest that mismatch responses may reflect signatures of Bayesian learning (Lieder 2013; Maheu 2019; Stefanics 2018; Ostwald 2012; Gijsen 2021). While studies directly investigating mismatch signals in response to multi-modal sensory inputs are rare, previous research indicates a ubiquitous role for cross-modal probabilistic learning. The brain tends to automatically integrate auditory, somatosensory and visual stimuli during sequence processing (Bresciani 2006; 2008; Frost 2015) and cross-modal perceptual associations can influence statistical learning of sequence regularities (Andric 2008; Parmentier 2011), modulate mismatch responses (Besle 2005; Butler 2012; Zhao 2015; Kiat 2018; Friedel 2020) and influence subsequent uni-modal processing in various ways (Shams 2011). Recent advances in modelling Bayesian causal inference suggest that the main computational stages of multi-modal inference evolve along a multisensory hierarchy involving early sensory segregation followed by mid-latency sensory fusion and late Bayesian causal inference (Rohe 2015; 2019; Cao 2019). However, the extent to which the MMN and P3 reflect these stages and should be considered sensory specific signatures of regularity violation or the result of modality independent computations in an underlying predictive network is not fully understood.

The current study aimed to investigate the commonalities and differences between mismatch responses in different modalities in a single experiment and to elucidate in how far they reflect local, uni-modal or global, cross-modal computations. To this end, we employed a roving stimulus paradigm, in which auditory, somatosensory and visual stimuli were simultaneously presented in a probabilistic tri-modal stimulus stream.

Typically, MMRs are studied with the oddball paradigm, in which rarely presented “oddball” stimuli deviate from frequently presented standard stimuli in some physical feature, such as sound pitch or stimulus intensity. The roving stimulus paradigm, on the other hand, defines deviants and standards in terms of their local sequence position, while the stimulus frequency (occurrence of stimulus types) across the sequence is equal (Cowan 1993; Baldeweg 2004). The deviant is defined as the first stimulus that breaks a train of repeating (standard) stimuli. With repetition, the deviant subsequently becomes the new standard, defining a train of stimulus repetitions. Thus, the roving stimulus paradigm is an excellent tool to experimentally induce mismatch responses, while controlling for differences in physical stimulus features.

Based on a probabilistic model, we generated sequences of high and low intensity stimuli that were governed by uni-modal transition probabilities as well as cross-modal conditional dependencies. This allowed us to test to what extent early and late mismatch responses are sensitive to local and global violations of statistical regularities and to draw conclusions regarding their potential role in cross-modal hierarchical inference. Specifically, we extracted the MMN and P3 MMRs for each modality and investigated their modality specific and modality general response properties regarding stimulus repetition and change, as well as their sensitivity to cross-modal predictive information. Further, we used source localization to investigate modality specific and modality general neuronal generators of mismatch responses. Finally, we complemented our average-based analyses with single-trial modelling to investigate if signatures of uni-modal and cross-modal Bayesian inference can account for trial-to-trial fluctuations in the MMN and P3 amplitudes.

## Materials and Methods

Participants underwent a novel multi-modal version of the roving stimulus paradigm. Our paradigm, depicted in figure 1, consisted of simultaneously presented auditory (A), somatosensory (S) and visual (V) stimuli, which each alternated between two different intensity levels (‘low’ and ‘high’). The tri-modal stimulus sequences originated from a single probabilistic model (described in the section *Probabilistic sequence generation*), resulting in different combinations of low and high stimuli across the three modalities in each trial.

**Figure 1:**
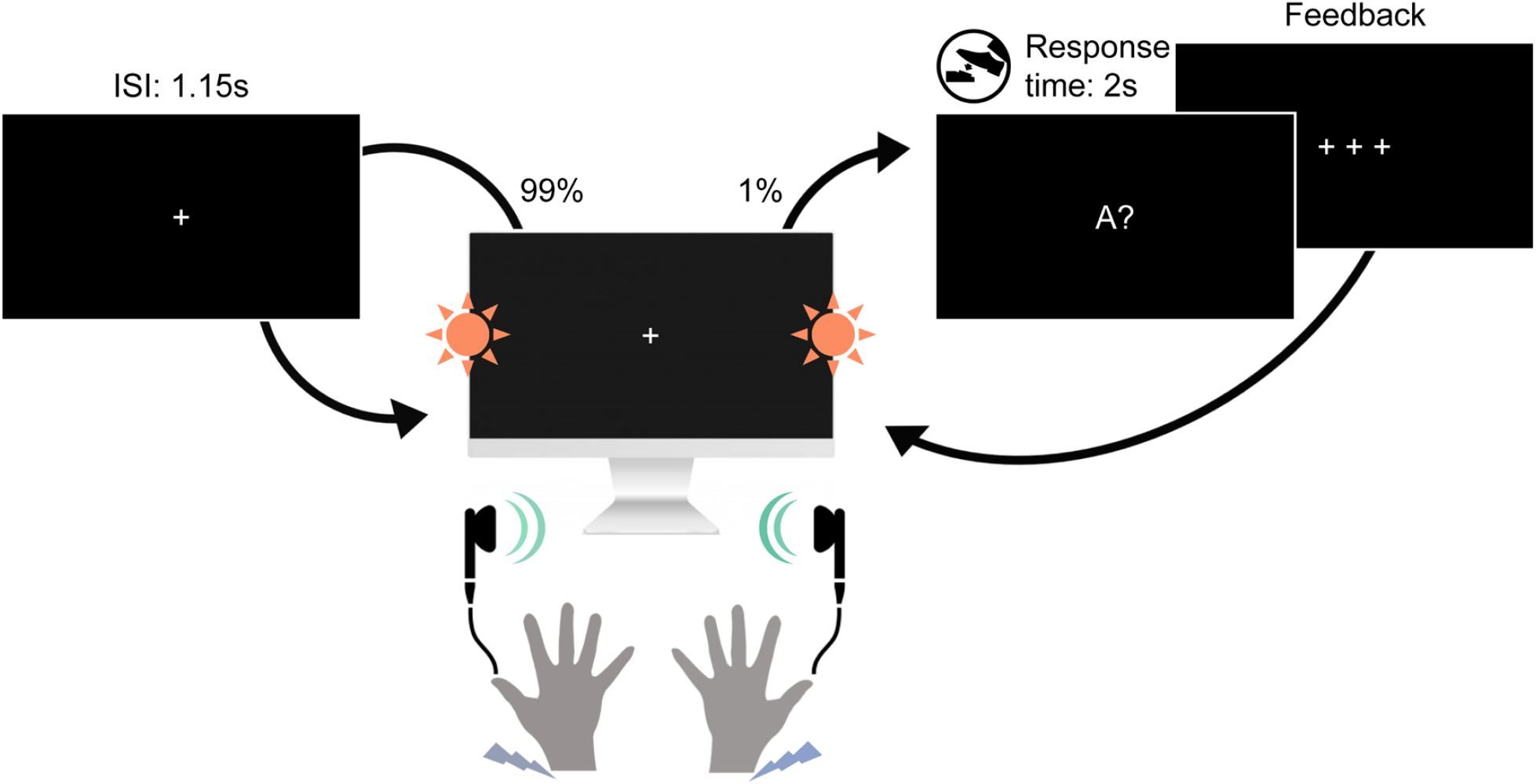
Experimental paradigm. Participants were seated in front of a screen and received sequences of simultaneously presented bilateral auditory beep stimuli (green), somatosensory electrical pulse stimuli (purple) and visual flash stimuli (orange) each at either low or high intensity. On consecutive trials, stimuli within each modality either repeated the previous stimulus intensity of that modality (standard) or alternated to the other intensity (deviant). This created tri-modal roving stimulus sequences, where the repetition/alternation probability in each modality was determined by a single probabilistic model (see *Probabilistic sequence generation*). In 1% of trials (catch trials) the fixation cross changed to one of the three letters A, T or V, interrupting the stimulus sequence. The letter prompted participants to indicate whether the last auditory (letter A), somatosensory (letter T for “tactile”) or visual (letter V) stimulus, respectively, was of high or low intensity. Responses were given with a left or right foot pedal press using the right foot.

### Participants

34 healthy volunteers (19-43 years old, mean age: 26, 22 females, all right-handed), recruited from the student body of the Freie Universität Berlin and the general public, participated for monetary compensation or an equivalent in course credit. The study was approved by the local ethics committee of the Freie Universität Berlin and written informed consent was obtained from all participants prior to the experiment.

### Experimental Setup

Each trial consisted of three bilateral stimuli (A, S and V) that were presented simultaneously by triggering three instantaneous outputs of a data acquisition card (National Instruments Corporation, Austin, Texas, USA) every 1150ms (inter-stimulus interval).

Auditory stimuli were presented via in-ear headphones (JBL, Los Angeles, California, USA) to both ears and consisted of sinusoidal waves of 500Hz and 100ms duration that were modulated by two different amplitudes. The amplitudes were individually adjusted with the participants to obtain two clearly distinguishable intensities (mean of the low intensity stimulus: 81.43 ± 1.22 *dB* ; mean of the high intensity stimulus: 93.02 ± 0.98 *dB*).

Somatosensory stimuli were administered with two DS5 isolated bipolar constant current stimulators (Digitimer Limited, Welwyn Garden City, Hertfordshire, UK) via adhesive electrodes (GVB-geliMED GmbH, Bad Segeberg, Germany) attached to the wrists of both arms. The stimuli consisted of electrical rectangular pulses of 0.2ms duration. To account for interpersonal differences in sensory thresholds, the two intensity levels used in the experiment were determined on an individual basis. The low intensity level (mean: 3.97 ± 0.84*mA*) was set in proximity to the detection threshold yet high enough to be clearly perceivable (and judged to be the same intensity on both sides). The high intensity level (mean: 6.47 ± 1.33*mA*) was determined for each participant to be easily distinguishable from the low intensity level yet remaining non-painful and below the motor threshold.

Visual stimuli were presented via light emitting diodes (LEDs) and transmitted through optical fiber cables mounted vertically centered to both sides of a monitor. The visual flashes consisted of rectangular waves of 100ms duration that were modulated by two different amplitudes (low intensity stimulus: 2.65 *V* ; high intensity stimulus: 10 *V*) that were determined to be clearly perceivable and distinguishable prior to the experiment. Participants were seated at a distance of about 60cm to the screen such that the LED’s were placed within the visual field at a visual angle of about 67 degrees.

In each of 6 experimental runs of 11.5 minutes, a sequence of 600 stimulus combinations was presented. To ensure that participants maintained attention throughout the experiment and to encourage monitoring of all three stimulation modalities, participants were instructed to respond to occasional catch trials (target questions) via foot pedals. In six trials randomly placed within each run the fixation cross changed to one of the letters A, T or V followed by a question mark. This prompted participants to report if the most recent stimulus (directly before appearance of the letter) in the auditory (letter A), somatosensory (letter T for “tactile”) or visual (letter V) modality was presented with low or high intensity. The right foot was used to press either a left or a right pedal, and the pedal assignment (left=low/right=high or left=high/right=low) was counterbalanced across participants.

### Probabilistic sequence generation

Each of the three sensory modalities (A, S, V) were presented as binary (low/high) stimulus sequences originating from a common probabilistic model. The model consists of a state *s* at time *t* evolving according to a Markov chain (*p*(*s*_*t*_|*s*_*t*−1_)) with each state deterministically emitting a combination of three binary observations conditional on the preceding observation combination (*p*(*o*_*A,t*_, *o*_*S,t*_, *o*_*V,t*_|*o*_*A,t*−1_, *o*_*S,t*−1_, *o*_*V,t*−1_)). For example, a transition expressed as [100|000] indicates a unimodal auditory change (*o*_*A,t*_ = 1, *o*_*A,t*−1_ = 0) with repeating somatosensory and visual modalities (*o*_*S,t*_ = *o*_*S,t*−1_ = 0 and *o*_*V,t*_ = *o*_*V,t*−1_ = 0). For each stimulus modality, in each state, the other two modalities form either congruent observations ([00] and [11]), or incongruent observations ([01] and [10]), which was used to manipulate the predictability of transitions in the sequences in different runs of the experiment.

Three types of stimulus sequences, depicted in figure 2 were generated with different probability settings. The settings determine the transition probabilities within each modality given the arrangement of the other two modalities (i.e. either congruent or incongruent). One setting defines lower change probability if the other two modalities are congruent (e.g. for any change in modality A from *t* − 1 to *t*, S and V were *congruent* with *p*(100|000) = *p*(000|100) = *p*(111|011) = *p*(011|111) = 0.025 and S and V were *incongruent* with *p*(101|001) = *p*(001|101) = *p*(110|010) = *p*(010|110) = 0.15). The second setting defines lower change probability if the other two modalities are incongruent (e.g. for any change in modality A from *t* − 1 to *t*, S and V were *incongruent* with *p*(101|001) = *p*(001|101) = *p*(110|010) = *p*(010|110) = 0.025 and S and V were *congruent* with *p*(100|000) = *p*(000|100) = *p*(111|011) = *p*(011|111) = 0.15). The third setting defines equal change probability if the other two modalities are congruent or incongruent (e.g. for any change in modality A from *t* − 1 to *t*, S and V were *congruent* with *p*(100|000) = *p*(000|100) = *p*(111|011) = *p*(011|111) = 0.0875 and S and V were *incongruent* with *p*(101|001) = *p*(001|101) = *p*(110|010) = *p*(010|110) = 0.0875).

**Figure 2:**
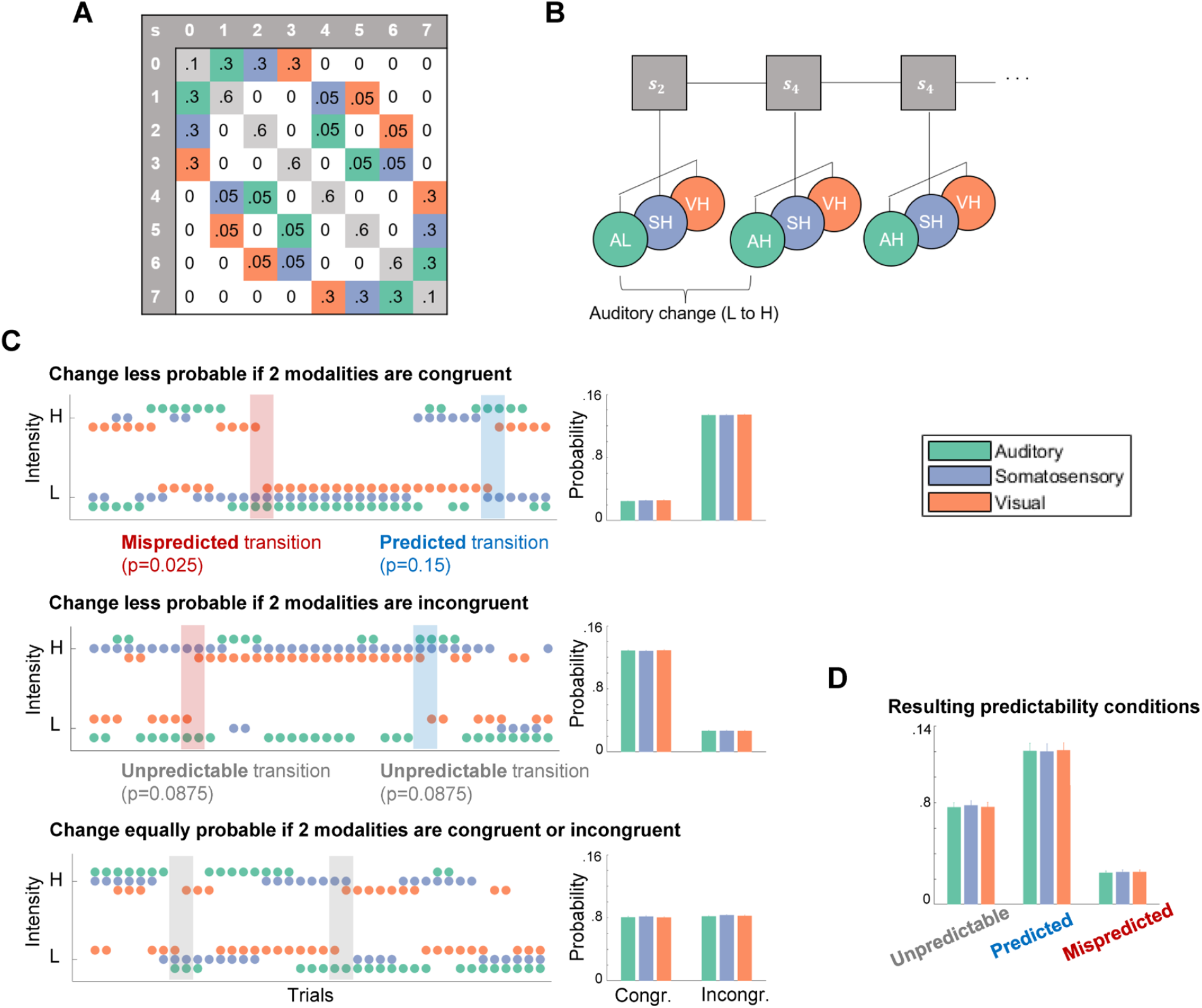
Probabilistic sequence generation. A) Schematic of state transition matrix. Colors depict transitions in the respective modality which were assigned specific transition probabilities: Green=auditory, purple=somatosensory, orange=visual, light-gray=tri-modal repetition, white=multi-modal change (set to zero). B) Visualization of states (s) evolving according to a Markov chain emitting tri-modal binary outcomes. C) Probability settings of stimulus sequences. Left column: Sequences. Right column: Averaged empirical change probabilities across all sequences. Top: Transition probabilities determine that for each modality a change is unlikely (p=0.025) if the other two modalities are congruent (and likely if they are incongruent; p=0.15). Middle: Transition probabilities determine that for each modality a change is likely (p=0.15) if the other two modalities are congruent (and unlikely if they are incongruent; p=0.025). Bottom: Transition probabilities determine that for each modality a change is equally likely (p=0.0875) if the other two modalities are congruent or incongruent. D) Averaged empirical change probabilities for predictability conditions.

In each of 6 experimental runs the stimulus sequence was defined by one of the three different probability settings. Each probability setting was used twice during the experiment and the order of the 6 different sequences was randomized. Participants were unaware of the sequence probabilities and any learning of sequence probabilities was considered to be implicit and task irrelevant.

Following the nomenclature suggested by Arnal and Giraud (2012), the resulting stimulus transitions for each modality within the different sequences can be defined as being either *predicted* (here higher change probability conditional on congruency/incongruency), *mispredicted* (here lower change probability conditional on congruency/incongruency) or *unpredictable* (here equal change probability). For each modality, repetitions are more likely (*p* = 0.825) than changes (*p* = 0.175) regardless of the type of probability setting and stimulus, resulting in classic roving standard sequences for each modality (mean stimulus train length: 5, mean range of train length: 2-34 stimuli).

### EEG data collection and preprocessing

Data was collected using a 64-channel active electrode EEG system (ActiveTwo, BioSemi, Amsterdam, Netherlands) at a sampling rate of 2048Hz, with head electrodes placed in accordance with the extended 10-20 system. Individual electrode positions were recorded using an electrode positioning system (zebris Medical GmbH, Isny, Germany).

Preprocessing of the EEG data was performed using SPM12 (Wellcome Trust Centre for Neuroimaging, Institute for Neurology, University College London, London, UK) and in-house MATLAB scripts (MathWorks, Natick, Massachusetts, USA). First, the data was referenced against the average reference, high-pass filtered (0.01Hz), and downsampled to 512Hz. Subsequently, eye-blinks were corrected using a topographical confound approach (Berg 1994; Ille 2002) and epoched using a peri-stimulus time interval of -100 to 1050ms. All trials were visually inspected and artefactual data removed. Likewise, catch trials were omitted for all further analyses. Furthermore, the EEG data of two consecutive participants were found to contain excessive noise due to hardware issues, resulting in their exclusion from further analyses and leaving data of 32 participants. Finally, a low-pass filter was applied (45Hz) and the preprocessed EEG data was baseline corrected with respect to the pre-stimulus interval of -100 to -5ms. To use the general linear model (GLM) implementation of SPM, the electrode data of each participant was linearly interpolated into a 32×32 grid for each time point, resulting in one three-dimensional image (with dimensions 32×32×590) per trial. These images were then spatially smoothed with a 12×12mm full-width half-maximum (FWHM) Gaussian kernel to meet the requirements of random field theory, which the SPM software uses to control the family wise error rate.

### Event-related responses and statistical analysis

First, to extract basic mismatch response signals of each modality from the EEG data, we contrasted standard and deviant trials of each modality with paired t-tests corrected for multiple comparisons by using cluster-based permutation tests implemented in fieldtrip (Maris 2007). Two time windows of interest were defined based on the literature (Duncan 2009) to search for earlier negative clusters between 50-300ms, corresponding to the mismatch negativity, and later positive clusters between 200-600ms, corresponding to the P3. Clusters were defined as adjacent electrodes with a cluster defining threshold of *p*_*fwe*_<0.05.

For further analyses, GLMs were set up as implemented in SPM12, which allows defining conditions on the single trial level. To test for effects of stimulus repetitions on standards, deviants and mismatch responses (deviants minus standards), a *TrainLength* model was defined that consisted of 45 regressors: an intercept regressor, 36 regressors coding for the repetition train length (trials binned into 1, 2, 3, 4-5, 6-8, >8 repetitions) for standards (i.e. the position of the standard in the current train) and deviants (i.e. the number of standards preceding the deviant) in each modality, as well as 4 global standard and 4 global deviant regressors. The global regressors captured the train length (1, 2, 3, >3 repetitions) of standards and deviants regardless of their modality, meaning that trials in which standards occurred in all three modalities were coded as global standards, whereas trials in which a deviant occurred in any of the three modalities were coded as global deviants.

To test for the implicit effect of cross-modal predictability based on the different conditional probability setting in the sequence, a *Predictability* model was defined that consisted of 37 regressors: an intercept regressor and 18 regressors coding standards and deviants of each modality for each of the three conditions described above: *unpredictable* (trials originate from sequences with no conditional dependence between modalities), *predicted* (trials originate from sequences with conditional dependence; trials defined by change being likely), *mispredicted* (trials originate from sequences with conditional dependence; trials defined by change being unlikely). On the single-participant level, these were coded for congruent and incongruent trials separately resulting in 36 regressors.

Finally, a *P3-Conjunction* model was specified that consisted of 7 regressors: an intercept regressor and 6 regressors coding all standards and deviants for each of the three modalities. This model was used to apply SPM’s second level conjunction analysis, contrasting standards and deviants across modalities in search of common P3 effects across modalities.

Each GLM was estimated on the single-trial data of each participant using restricted maximum likelihood estimation. This yielded *β*-parameter estimates for each model regressor over (scalp-) space and time, which were subsequently analyzed at the group level. Second level analyses consisted of a mass-univariate multiple regression analysis of the individual *β* scalp-time images with a design matrix specifying regressors for each condition of interest as well as a subject factor. Second level beta estimates were contrasted for statistical inference and multiple comparison correction was achieved with SPM’s random field theory-based FWE correction (Kilner 2005).

### Source localization

To investigate the most likely underlying neuronal sources for the mismatch negativity and P3 mismatch response we applied distributed source reconstruction as implemented in SPM12 to the ERP data. For each participant, the MMN of each modality (auditory, somatosensory, visual) was source localized within a time window of 100-200ms. For the P3, the average MMR at 330ms was chosen for source localization as this time point showed the strongest overlap of P3 responses between modalities (based on the results of the P3 conjunction contrast).

Participant-specific forward models were created using an 8196-vertex template cortical mesh co-registered with the individual electrode positions via fiducial markers. An EEG head model based on the boundary element method (BEM) was used to construct the forward model’s lead field. For the participant-specific source estimates, multiple sparse priors under group constraints were applied. The source estimates were subsequently analyzed at the group level using the GLM implemented in SPM12. Second-level contrasts consisted of one-sample t-tests for each modality as well as (global) conjunction contrasts across modalities. The resulting statistical parametric maps were thresholded at the peak level with p<0.05 after FWE correction. The anatomical correspondence of the MNI coordinates of the cluster peaks were identified via cytoarchitectonic references using the SPM Anatomy toolbox (Eickhoff 2005).

### Single-trial modelling of EEG data

In addition to the analysis of event-related potentials, the study aimed to compare different computational strategies of sequence processing potentially underlying neuronal generation of mismatch responses. To this end, we generated regressors from different Bayesian learning (BL) models as well as a train length dependent change detection (TLCD) model making different predictions for the single-trial EEG data.

Theories on MMN generation hypothesize adaptation and memory-trace dependent change detection to contribute to the MMN. With prior repetition of stimuli, the response to standard stimuli tends to decrease while the response to deviant stimuli tends to increase. We defined the TLCD model to reflect such reciprocal dynamics of responses to stimulus repetition and change without invoking assumptions of probabilistic inference. The model is defined for each modality separately and tracks the stimulus train lengths *c* for a given modality by counting stimulus repetitions: *c*_*t*_ = *d*_*t*_(*c*_*t*−1_ + *d*_*t*_), where 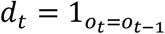 takes on the value 1 whenever the current observation *o*_*t*_ is a repetition of the previous observation *o*_*t*−1_ and *d*_*t*_ = 0 resets the current train length to zero. To form single-trial predictors of the EEG data, the model outputs values that increase linearly with train length and have opposite signs for standards and deviants:

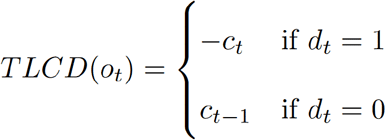

In addition to the TLCD model, different BL models were created to contrast the static train length based TLCD model with dynamic generative models tracking transition probabilities. The BL models consist of conjugate Dirichlet-Categorical models estimating probabilities of observations read out by three different surprise functions: Bayesian surprise (BS), Predictive surprise (PS) and confidence-corrected surprise (CS).

BS quantifies the degree to which an observer adapts their generative model to incorporate new observations (Itti 2009; Baldi 2010) and is defined as the Kullback-Leibler (KL) divergence between the belief distribution prior and posterior to the update: *BS*(*y*_*t*_) = *KL*(*p*(*s*_*t*−1_|*y*_*t*−1_, …, *y*_1_)||*p*(*s*_*t*_|*y*_*t*_, …, *y*_1_)). PS is based on Shannon’s (1948) definition of information and defined as the negative logarithm of the posterior predictive distribution, assigning high surprise to observed events *y*_*t*_ with low estimated probability of occurrence: *PS*(*y*_*t*_) = − *ln p*(*y*_*t*_|*s*_*t*_) = − *ln p*(*y*_*t*_|*y*_*t*−1_, …, *y*_1_). CS additionally considers the commitment of the generative model and scales with the negative entropy of the prior distribution (Faraji 2018). It is defined as the KL divergence between the (informed) prior distribution at the current time step and a flat prior distribution 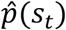 updated with the most recent event 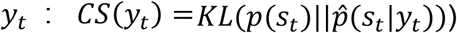.

Following Faraji et al. (2018) surprise quantifications can be categorized as puzzlement or enlightenment surprise. While puzzlement refers to the mismatch between sensory input and internal model belief, closely related to the concept of prediction error, enlightenment refers to the update of beliefs to incorporate new sensory input. In the current study we were interested in a quantification of the model inadequacy by means of an unsigned prediction error as reflected by surprise. As such, throughout the manuscript, with prediction error we do not refer to the specific term of (signed) reward prediction error as used for example in reinforcement learning but rather use it to refer to the signaling of prediction mismatch. While both PS and CS are instances of puzzlement surprise, CS is additionally scaled by belief commitment and quantifies the concept that a low-probability event is more surprising if commitment to the belief (of this estimate) is high. BS, on the other hand, is an instance of enlightenment surprise and is considered a measure of the update to the generative model resulting from new incoming observations.

A detailed description of the Bayesian observer, its transition probability version as well as the surprise read-out functions can be found in our previous work on somatosensory mismatch responses (Gijsen 2021). Here, we will primarily provide a brief description of the specifics of two implementations of Dirichlet-Categorical observer models, a uni- and a cross-modal model. Both observer models receive stimulus sequences (of one respective modality) as input and iteratively update a set of parameters with each new incoming observation. In each iteration the estimated parameters are read out by the surprise functions (BS, PS and CS) to produce an output which is subsequently used as a predictor for the EEG data.

For each modality, the uni-modal Dirichlet-Categorical model considers a binary sequence with two possible stimulus identities (low and high) estimating transition probabilities with *y*_*t*_ = *o*_*t*_ for *t* = 1, …, *T* with a set of hidden parameters *s*^(*i*)^ for each possible transition from *o*_*t*−1_ = *i*. This uni-modal model does not capture any cross-modal dependencies in the sequence (i.e. the alternation and repetition probabilities conditional on the tri-modal stimulus configuration). Therefore, we defined a cross-modal Dirichlet-Categorical model to address the question whether the conditional dependencies were used by the brain during sequence processing for prediction of stimulus change. The dependencies in the sequence were independent of stimulus identity but provide information about the probability of repetition or alternation (*d*_*t*_) conditional on the congruency of the other modalities. The cross-modal model thus estimates alternation probabilities (*y*_*t*_ = *d*_*t*_ for *t* = 2, …, *T*) with a set of hidden parameters *s*^(*i*)^ when other modalities are incongruent and *s*^(*c*)^ when other modalities are congruent. Therefore, while the uni-modal model learns the probability of stimulus *transitions* within modality, the cross-modal model learns the probability of stimulus *alternations* within modality conditional on the congruency of the other modalities. As such, the cross-modal model provides a minimal implementation of a Bayesian observer that captures the cross-modal dependencies in the sequences.

#### Model fitting procedure

The technical details of the model fitting and subsequent Bayesian model selection procedures are identical to Gijsen et al. (2021) where the interested reader is kindly referred to for further information. First, the stimulus sequence-specific regressor of each model was obtained for each participant. After z-score normalization, the regressors were fitted to the single-trial, event-related electrode data using a free-form variational inference algorithm for multiple linear regression (Flandin 2007; Penny 2005; Penny 2003). The obtained model-evidence maps were subsequently subjected to the Bayesian model selection (BMS) procedure implemented in SPM12 (Stephan 2009) to draw inferences across participants with well-established handling of the accuracy-complexity trade-off (Woolrich 2012).

In total, 8 regression models were fit: A null model (offset only), a TLCD regression model and, for each of the three surprise read-out functions, one regression model including only the uni-modal regressors and one additionally including the cross-modal regressors. The purely uni-modal regression model will be called UM and the regression model including uni-modal and, additionally, cross-modal regressors will be called UCM. The design matrix of the TLCD regression model consisted of 4 regressors, an offset and the predicted parametric change responses for each of the three modality sequences (auditory, somatosensory, visual). Similarly, the design matrix of the UM regression model consisted of 4 regressors, an offset and the surprise responses of the uni-modal model for each of the three modalities. The UCM regression model was identical to the UM regression model but with an additional three regressors containing the cross-modal surprise responses for each modality. Therefore, the UCM regression model is more complex and only gets assigned higher model evidence than the reduced UM regression model only if the additional regressors contribute significantly to a better model fit (Stephan 2009).

To allow for the possibility of different timescales of stimulus integration (Maheu 2019; Ossmy 2013; Runyan 2017), the integration parameter *τ* of the Dirichlet-Categorical model was optimised for each model, participant and peri-stimulus time-bin before model selection. To this end, model regressors were fit for a range of 11 tau parameter configurations ([0, 0.001, 0.0015, 0.002, 0.003, 0.005, 0.01, 0.02, 0.05, 0.1, 0.2]) corresponding to integration windows with a 0.5 stimulus weighting at (half-life of) [600, 462, 346, 231, 138, 69, 34, 13, 6, 3] stimuli, of which the parameter with the best model-evidence was chosen.

#### Bayesian model comparison

The estimated model-evidence maps were used to evaluate the models’ relative performance across participants via family-wise Bayesian model selection (Penny 2010). The model space was partitioned into three types of families to draw inference on different aspects of the involved models. Given that the literature provides some evidence for each of the three surprise read-out functions (BS, PS, CS) to capture some aspect of EEG mismatch responses, we included all of them in the family wise comparisons to avoid biasing the comparison of different BL models.

The first model comparison considered the full space of Bayesian learning models as a single family (BL family) and compared it to the TLCD model (TLCD family) and the null model (NULL family). Since the BL models had their tau parameter optimized, which was not possible for the TLCD model, we applied the same penalization method used in our previous study (Gijsen 2021). The degree to which the optimization on average inflated model evidence was subtracted from the BL models prior to BMS. Specifically, for all parameter values, the difference between the average model evidence and that of the optimized parameter was computed and averaged across post-stimulus time bins, electrodes and participants.

Subsequent analyses grouped the different BL models into separate families: The second comparison grouped the BL models into two families of uni-modal (UM) and cross-modal (UCM) models, as well as the null model, to test which electrodes and time points showed influences of uni-versus multi-modal processing. The third comparison grouped the BL models into three surprise families and the null model, to test whether the observed MMRs were best captured by predictive surprise (PS), Bayesian surprise (BS) or confidence-corrected surprise (CS).

## Results

### Behavioral results

Participants showed consistent performance in responding to the catch trials during each experimental run, indicating their ability to globally maintain their attention to the tri-modal stimulus stream. Of the 85.5% responses made in time, 75.3% were correct with an average reaction time of 1.4 ± 0.25*s*.

### Event-related potentials

#### Uni-modal mismatch responses

Cluster based permutation tests confirmed the presence of early modality specific MMN components as well as later P3 MMRs for all three modalities. Both early and late MMRs showed a modulation by the number of stimulus repetitions, the details of which will be described in the following sections.

#### Auditory MMRs

The MMN, as the classic mismatch response, has originally been studied in the auditory modality and is commonly described as the ERP difference wave calculated by subtraction of standard trials from deviant trials (deviants-standards). This difference wave typically shows a negative deflection at fronto-central electrodes and corresponding positivity at temporo-parietal sites, ranging from around 100-250ms (Näätänen 1978; Näätänen 1979; Näätänen 2007). Correspondingly, we find a significant negative fronto-central auditory MMN cluster between 80-200ms (figure 3A). Within the MMN cluster, deviants appear to deflect from the standard ERP around the peak of the auditory N1 component and reach their maximum difference around the peak of the subsequent P2 component. In the later time window, we observe positive MMRs at central electrodes between 200-400ms, corresponding to a P3 modulation, as well as beyond 400ms at progressively more posterior electrodes.

**Figure 3.**
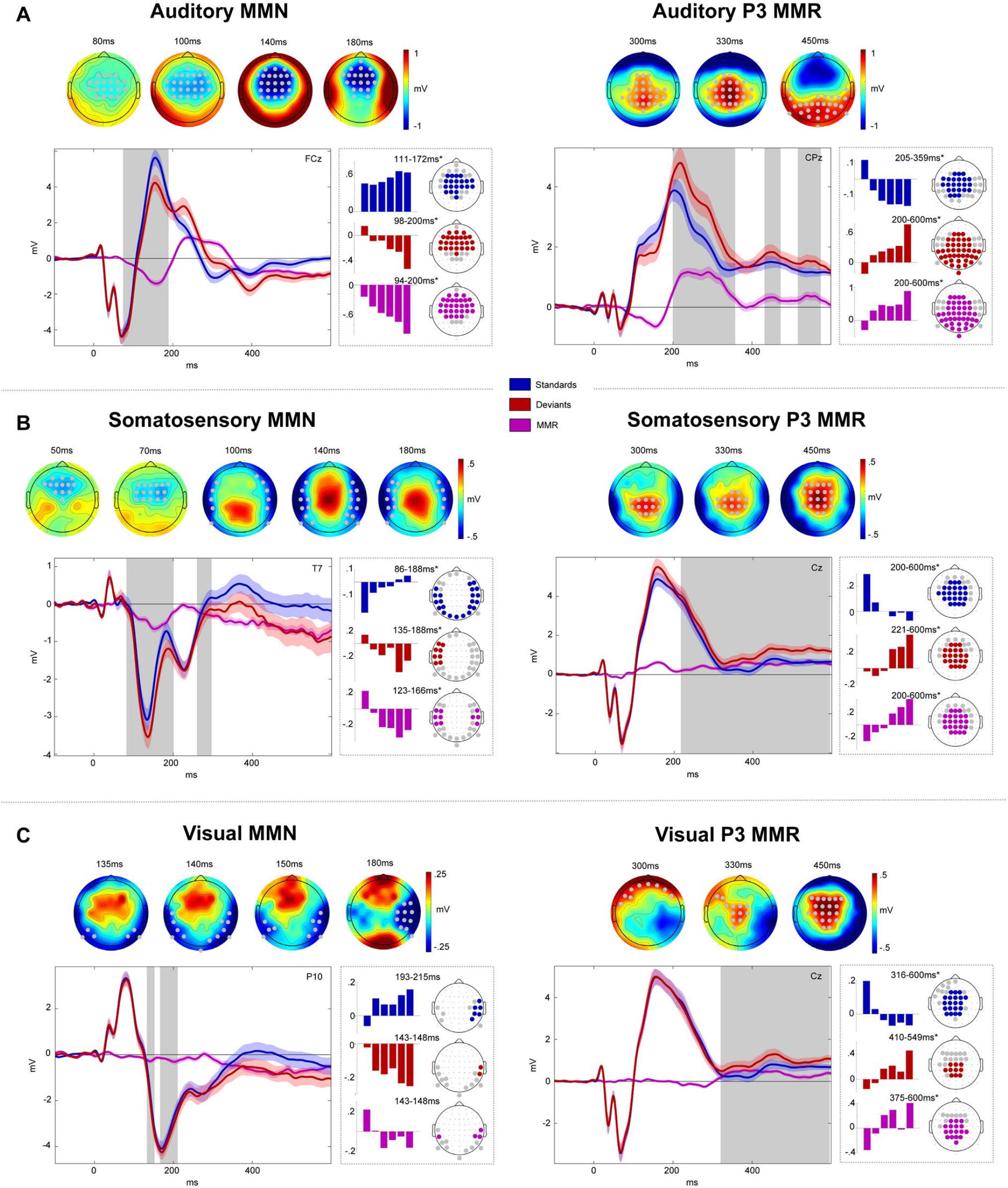
Mismatch responses. Panels A-C show MMRs of auditory (A), somatosensory (B) and visual (C) modalities. Within panels: Left: MMN. Right: P3 MMR. Gray dots (top) and gray boxes (bottom) indicate significant MMR electrodes and time points with *p*_*fwe*_ <0.05. Top row: MMR scalp topographies (deviants-standards). Bottom row: Grand average ERPs (left panels) and beta parameter estimates of significant linear contrast clusters (right panels). Colored bars depict six beta parameter estimates of the *TrainLength* GLM (1, 2, 3, 4-5, 6-8, >8 repetitions) averaged across electrodes within linear contrast clusters. Asterisks indicate significance of the linear contrast (*p*_*fwe*_ <0.05).

Within early and late auditory MMR clusters, the response to both standards and deviants was modulated by the number of standard repetitions. The auditory system is known to be sensitive to stimulus repetitions, particularly within the roving standard paradigm (Ulanovsky 2003; 2004; Cowan 1993; Baldeweg 2004). Therefore, we hypothesised a gradual increase of the auditory response to standard stimuli around the time of the MMN, known as repetition positivity (Baldeweg 2004; Baldeweg 2006; Haenschel 2005) as well as reciprocal negative modulation of the corresponding deviant response (Näätänen 2007; Bendixen 2007). Together, these effects should result in a gradual increase of the MMN amplitude with stimulus repetition. Indeed, linear contrasts applied to the GLM beta parameter estimates of the *TrainLength* model revealed that the MMN increases with the repetition of standards before a deviant was presented (94-200ms, cluster *p*_*f*we_<0.001). This effect was driven by a negative linear modulation of the deviant response (98-200ms, cluster *p*_*f*we_ <0.001) as well as a repetition positivity effect on the standards (111-172ms, cluster *p*_*f*we_ <0.001). Similarly, the later P3 MMR increased with standard repetitions (200-600ms, cluster *p*_*fwe*_ <0.001) and this effect was driven by an increase of deviant responses (200-600ms, cluster *p*_*fwe*_ <0.001) and a decrease of standard responses (205-359ms, cluster *p*_*fwe*_<0.001). Given the temporal difference between standard (around 200-350ms) and deviant (200-600ms) train length effects, the parametric modulation of the late MMR beyond 350ms seems to be primarily driven by the increase in deviant responses.

#### Somatosensory MMRs

We hypothesized somatosensory MMRs to consist of early bilateral (fronto-) temporal negativities, resulting primarily from increased N140 components (Kekoni 1997), with a corresponding central positivity extending into a later central P3 component.

After an early mismatch effect starting at ∼50ms at fronto-central electrodes, a more pronounced bilateral temporal cluster emerged that extended from ∼90-190ms and can be considered the somatosensory equivalent of the auditory MMN (figure 3B). A reversed positive central component can be observed at the time of the somatosensory MMN (sMMN) and throughout the entire later time window (200-600ms) at which point it can be considered a putative P3 MMR.

Early and late somatosensory MMRs were significantly modulated by stimulus repetition. Bilateral electrodes within the sMMN cluster show an increase of the sMMN amplitude with repetition (123-166ms, cluster *p*_*fwe*_<0.05). This effect was driven by an increase of deviant negativity (135-188ms, cluster *p*_*fwe*_<0.05) in combination with a positivisation of the standard (86-188ms, cluster *p*_*fwe*_ <0.05). Similarly, the later P3 MMR increases with repetition of standards (200-600ms, cluster *p*_*fwe*_<0.05), mutually driven by increasing deviant responses (221-600ms, cluster *p*_*fwe*_ <0.05) and decreasing standard responses (200-600ms, cluster *p*_*fwe*_<0.05).

#### Visual MMRs

We hypothesized visual MMRs to present as an early MMN at occipital to parieto-temporal electrodes and a later P3 component at central electrodes. Although less pronounced than its auditory and somatosensory counterparts, we indeed observed a negative visual mismatch component that developed from occipital to parieto-temporal electrodes between ∼130-200ms (figure 3C). In the later time window, we found a central positive component between ∼300-600ms, corresponding to a P3 MMR.

Within the significant visual MMN (vMMN) cluster, the linear contrast testing for repetition effects did not reach significance when correcting clusters for multiple comparisons (*p*_*fwe*_ >0.05). However, it is worth noting that some electrodes in this cluster seemed to show a similar pattern of response increases and decreases as in the auditory and somatosensory modality, which became apparent at more lenient thresholds. The vMMN tended to become more negative with repetition of standards (143-148ms, peak *p*_*uncorr*_ <0.005), with opposite tendencies of deviant negative increase (143-148ms, peak *p*_*uncorr*_<0.05) and standard decrease (193-215ms, peak *p*_*fwe*_<0.05). Thus, although we cannot conclude a modulation by standard repetition of the vMMN with any certainty, the observed beta parameters are in principle compatible with the effects observed in the auditory and somatosensory modalities (please see the discussion for potential reasons for the reduced vMMN in our data).

Within the P3 MMR cluster, on the other hand, we find significant clusters of linear increase of the MMR (375-600ms, cluster *p*_*fwe*_<0.05), again constituted by an increase in deviant responses (410-549ms, cluster *p*_*fwe*_ <0.05) and concomitant decrease in standard responses (316-600ms, cluster *p*_*fwe*_<0.05).

#### Cross-modal P3 effects

In search of a common P3 effect to deviant stimuli, we created conjunctions of the *deviants>standards* contrasts across the auditory, somatosensory and visual modalities. The conjunction revealed a common significant cluster starting at ∼300ms (cluster *p*_*fwe*_<0.05) that comprised anterior central effects around 300-350ms followed by more posterior effects from 400-600ms (figure 4A).

**Figure 4.**
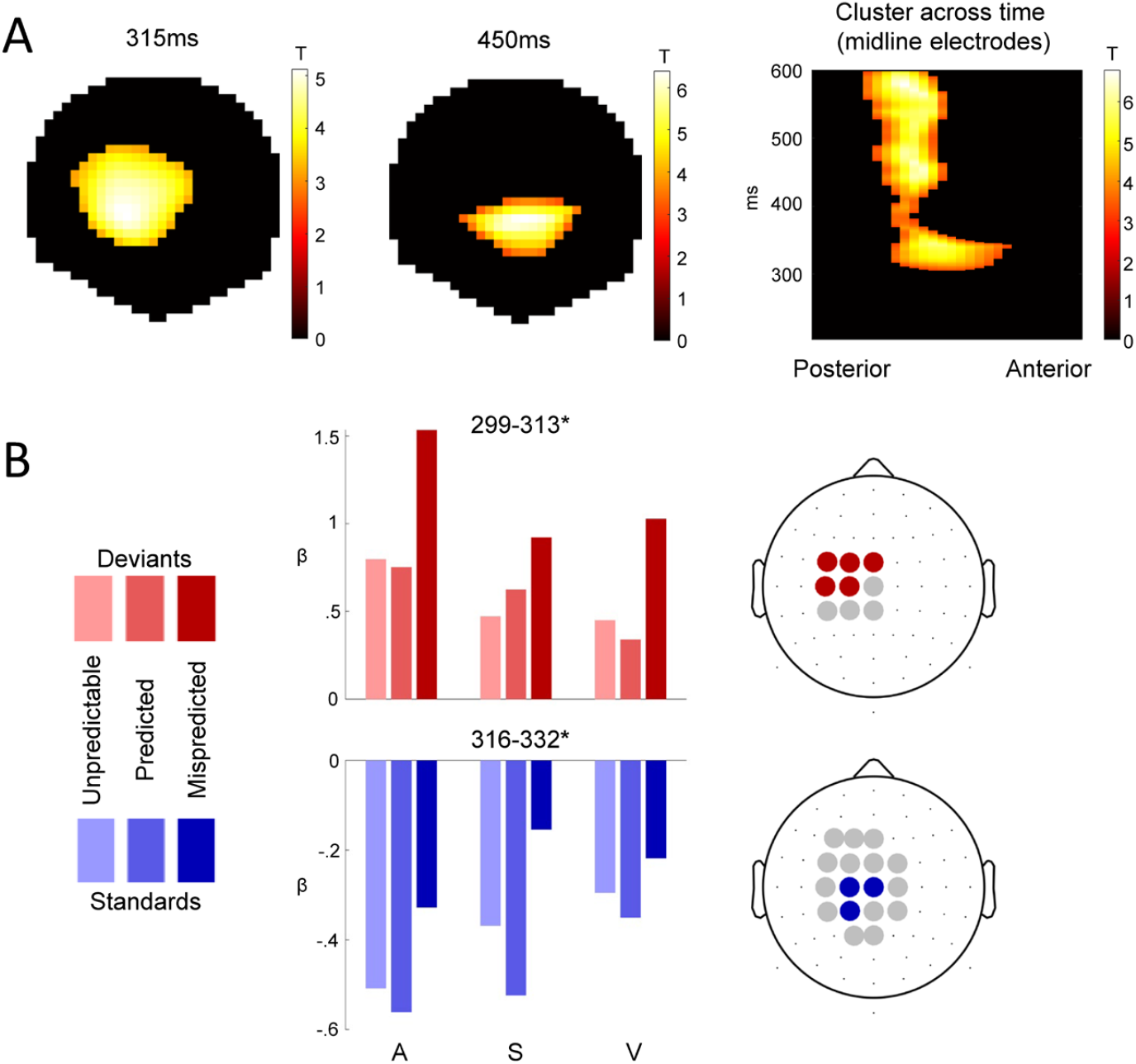
Cross-modal P3 effects. A) T-Maps of the conjunction of deviant>standard contrasts across the auditory, somatosensory and visual modalities. B) Beta estimates averaged across electrodes within significant clusters with peak *p*_*fwe*_ <0.05, resulting from two-way ANOVAs testing for differences between *unpredictable, predicted* and *mispredicted* deviants (red) and standards (blue).

To investigate the modulation of the P3 MMR by predictability we used two-way ANOVAs with the three-level factor *modality* (*auditory, somatosensory, visual*) and the three-level factor *predictability* condition (*predicted, mispredicted, unpredictable*). Separate ANOVAs were applied to deviants and standards. We hypothesized that the cross-modal P3 MMR might be sensitive to multisensory predictive information in the sequence, as the P3 has been shown to be sensitive to global sequence statistics (Wancongne 2011; Bekinschtein 2009) and to be modulated by stimulus predictability (Ritter 1999; Sussman 2003; Horvath 2008; Horvath 2012; Max 2015; Prete 2022). Indeed, within the common P3 cluster, both deviants (299-313ms, peak *p*_*fwe*_<0.05) and standards (316-332ms, peak *p*_*fwe*_ <0.05) show significant differences between predictability conditions. No significant interaction of predictability condition with modality was observed.

Post-hoc t-tests were applied to the peak beta estimates to investigate the differences between the three pairs of conditions. For the ANOVA concerning the deviant trials, post-hoc t-tests show a significant difference for *mispredicted>predicted* (t=14.667; p<0.001, Bonferroni corrected), *mispredicted>unpredictable* (t=14.76; p<0.001, Bonferroni corrected) and no significant difference between *unpredictable>predicted* conditions (t=0.01; p>0.05). Similarly, for the ANOVA concerning the standard trials, post-hoc t-tests show that there is a significant difference for *mispredicted>predicted* (t=10.67; p<0.001, Bonferroni corrected), *mispredicted>unpredictable* (t=6.87; p<0.001, Bonferroni corrected) and *unpredictable>predicted* conditions (t=3.83; p<0.001, Bonferroni corrected).

Taken together, this result suggests that stimuli which were mispredicted based on the predictive multisensory configuration resulted in increased responses within the common P3 cluster compared to predicted or unpredictable stimuli, regardless of their role as standards or deviants in the current stimulus train.

For completeness, we also tested the effect of predictability in the earlier MMN cluster, but we did not observe any significant modulations here (results not shown).

### Source localization

The source reconstruction analysis resulted in significant clusters of activation for each modality’s MMN as well as the P3 MMR. The results are depicted in figure 5 and cytoarchitectonic references are described in table 1.

**Table 1.** Source localization and cytoarchitectonic reference.

| Contrast | hemi-sphere | cytoarchitecture (probability) | MNI coord. at cytoarch. | t statistics at cytoarch. (p-value) |
| --- | --- | --- | --- | --- |
| aMMN | left | Auditory areas:<br>TE 4 (67.3%)<br>TE 3 (15.1%)<br>TE 1 (50.7%) | -52 -26 0<br>-59.8 -17.6 5.4<br>-50.3 -19.2 5.8 | 6.2 ($p_{fwe} < 0.05$)<br>4.81 ($p_{fwe} < 0.05$)<br>4.05 ($p_{uncorr} < 0.001$) |
| aMMN | right | Auditory areas:<br>TE 4 (54.1%)<br>TE 3 (33.6%)<br>TE 1 (61.9%) | -56 -26 0<br>-64.2 -16.4 5<br>53 -10.1 3.8 | 7.64 ($p_{fwe} < 0.05$)<br>5.85 ($p_{fwe} < 0.05$)<br>3.53 ($p_{uncorr} < 0.001$) |
| sMMN | left | Somatosensory areas:<br>3b [S1] (31.4%)<br>OP4 [S2] (38.6%)<br>OP1 [S2] (16%) | -15.7 -33.7 68.6<br>-65 -14.8 20.1 | 4.8 ($p_{fwe} < 0.05$)<br>3.72 ($p_{uncorr} < 0.001$) |
| sMMN | right | Somatosensory areas:<br>3b [S1] (40.3%)<br>OP4 [S2] (prob. 46.9%)<br>OP1 [S2] (prob. 9.1%) | 13.3 -33.7 68.2<br>66.5 -10.6 20.9 | 4.76 ($p_{fwe} < 0.05$)<br>3.51 ($p_{uncorr} < 0.001$) |
| vMMN | left | Visual areas:<br>hOc1 [V1] (84.6%)<br>hOc2 [V2] (11.1%)<br>hOc4v [V4] (51%)<br>hOc3v [V3] (24.6%)<br>FG4 (89.5%) | -10 -100 0<br>-30 -87.9 -11.7<br>-43.5 -49.4 -13.3 | 6.18 ($p_{fwe} < 0.05$)<br>5.51 ($p_{fwe} < 0.05$)<br>3.83 ($p_{uncorr} < 0.001$) |
| vMMN | right | Visual areas:<br>hOc1 [V1] (86.4%)<br>hOc2 [V2] (11%)<br>hOc3 [V3] (43.7%)<br>hOc4 [V4] (23.8%)<br>FG4 (59.8%) | 20 -100 -4<br>34.4 -88.7 -7.7<br>50.3 -42.5 -18.8 | 5.17 ($p_{fwe} < 0.05$)<br>4.7 ($p_{fwe} < 0.05$)<br>3.82 ($p_{uncorr} < 0.001$) |
| MMN<br>con-<br>junction | left | Frontal pole (28%)<br>Inferior temporal gyrus (49%) | -14 62 8<br>-50 -24 -28 | 2.56 ($p_{fwe} < 0.05$)<br>2.52 ($p_{fwe} < 0.05$) |
| MMN<br>con-<br>junction | right | Middle frontal gyrus (36%)<br>Inferior frontal gyrus (53%)<br>Frontal pole (65%)<br>Inferior parietal lobe (46.8%) | 42 28 30<br>53 27.6 17.3<br>30 42 32<br>62 -14 24 | 2.89 ($p_{fwe} < 0.05$)<br>2.96 ($p_{fwe} < 0.05$)<br>2.28 ($p_{fwe} < 0.05$)<br>2.85 ($p_{fwe} < 0.05$) |
| P3<br>con-<br>junction | (left) | Anterior cingulate gyrus<br>(51%) | -6 14 36 | 4.34 ($p_{fwe} < 0.05$) |
| P3<br>con-<br>junction | left | Frontal pole (74%)<br>Inferior frontal gyrus (35%)<br>Middle frontal gyrus (78%)<br>Superior frontal gyrus (43%)<br>Inferior temporal gyrus (34%)<br>Lateral occipital [hOc4la] (81%) | -34 44 20<br>52 26 18<br>-44 30 32<br>-22 12 60<br>-48 -44 -24<br>-46 -80 -8 | 3.31 ($p_{fwe} < 0.05$)<br>2.74 ($p_{fwe} < 0.05$)<br>2.87 ($p_{fwe} < 0.05$)<br>3.13 ($p_{fwe} < 0.05$)<br>3.21 ($p_{fwe} < 0.05$)<br>2.99 ($p_{fwe} < 0.05$) |
| P3<br>con-<br>junction | right | Frontal pole (86%) | 45 42 10 | 3.57 ($p_{fwe} < 0.05$) |
| | | Inferior frontal gyrus (39.5%) | 52 26 18 | 3.45 ($p_{fwe} < 0.05$) |
| | | Middle frontal gyrus (38%) | 36 2 54 | 3.0 ($p_{fwe} < 0.05$) |
| | | Precentral gyrus [4a] (19%) | 6 -32 64 | 2.64 ($p_{fwe} < 0.05$) |
| | | Lateral occipital cortex [hOc5] (55%) | 56 -62 0 | 3.45 ($p_{fwe} < 0.05$) |

**Figure 5.**
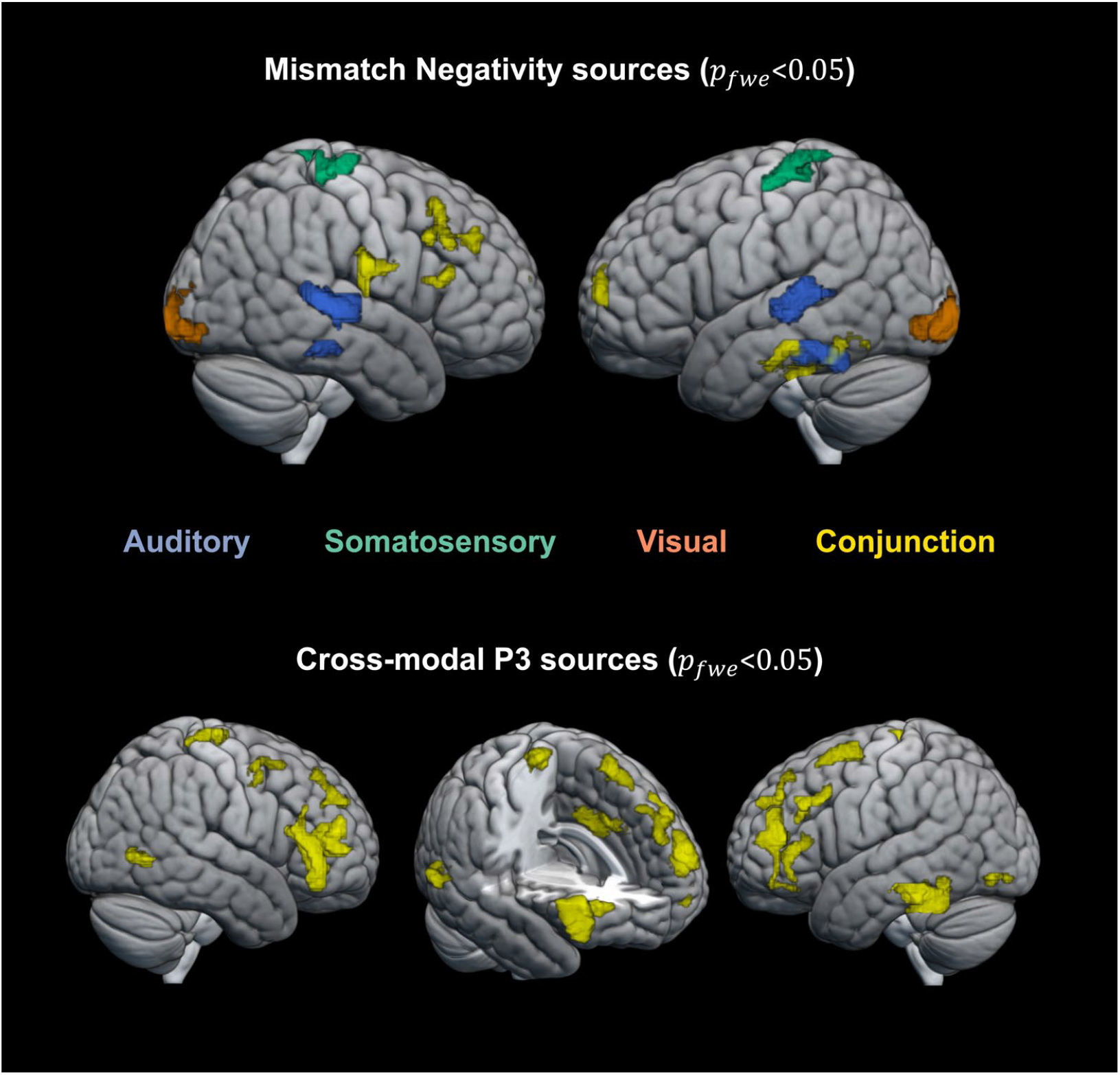
Source localisation. Top row: significant sources (*p*_*fwe*_ <0.05) for the auditory (purple), somatosensory (green) and visual (orange) MMNs as well as their conjunction (yellow). Bottom row: significant sources (*p*_*fwe*_ <0.05) for the conjunction (yellow) of the P3 MMR in the auditory, somatosensory and visual modalities.

For each modality, the MMN was localized to source activations in the respective modality’s sensory cortex and frontal cortex. Source localization of the auditory MMN shows the strongest activation in bilateral superior temporal areas (*p*_*fwe*_ <0.05; left cluster: peak t=6.20; right cluster: peak t=7.64) corresponding to auditory cortex and in inferior temporal areas (*p*_*fwe*_<0.05; left cluster: peak t=5.63; right cluster: peak t=5.60). The somatosensory MMN shows highest source activation in postcentral gyrus (*p*_*fwe*_ <0.05; left cluster: peak t=5.22; right cluster: peak t=4.92) corresponding to primary somatosensory cortex. Similarly, the visual MMN shows highest source activation in the occipital cortex (*p*_*fwe*_<0.05; left cluster: peak t=6.18; right cluster: peak t=5.17), around the occipital pole, corresponding to visual areas (V1-V4). Lowering the threshold to *p*_*uncorr*_ <0.001 (only shown in table 1) suggests additional activation of hierarchically higher sensory areas such as secondary somatosensory cortex for the sMMN (*p*_*uncorr*_ <0.001; left cluster: peak t=4.21; right cluster: peak t=5.01) and lateral occipital cortex (fusiform gyrus) for vMMN (part of the primary visual cluster). In addition to the sensory regions, common frontal sources with dominance on the right hemisphere were identified using a conjunction analysis for the MMN of all three modalities. In particular, significant common source activations were found in the right inferior frontal gyrus (*p*_*fwe*_<0.05; cluster: peak t=3.15) and right middle frontal gyrus (*p*_*fwe*_0.05; cluster: peak t=2.89). Additional significant common sources include frontal pole (*p*_*fwe*_<0.05; left cluster: peak t=2.56; right cluster: peak t=2.28), left inferior temporal gyrus (*p*_*fwe*_<0.05; cluster: peak t=2.52) and right inferior parietal lobe (*p*_*fwe*_<0.05; cluster: peak t=2.85).

For the late P3 MMR a wide range of sources was expected to contribute to the EEG signal (Linden 2005; Sabeti 2016). To identify those that underlie the P3 MMR common to all modalities, we used a conjunction analysis. Significant clusters were found primarily in anterior cingulate cortex (*p*_*fwe*_ <0.05; cluster: peak t=4.34) and bilateral (pre-)frontal cortex (*p*_*fwe*_<0.05; left inferior frontal gyrus cluster: peak t=3.57; left superior frontal gyrus cluster: peak t=3.13; left middle frontal gyrus cluster: peak t=2.87; left frontal pole cluster: peak t=3.31; right inferior frontal gyrus cluster: peak t=3.45; right middle frontal gyrus cluster: peak t=3.0; right frontal pole cluster: peak t=3.57). Additional significant sources were found in left inferior temporal gyrus (*p*_*fwe*_<0.05; cluster: peak t=3.21), left and right lateral occipital cortex (*p*_*fwe*_<0.05; left cluster: peak t=2.99; right cluster: peak t=3.45) and right precentral gyrus (*p*_*fwe*_<0.05; cluster: peak t=2.64).

### Single-trial modelling

As described in the previous sections, the responses of standards and deviants show specific sensitivity to (1) stimulus repetition and (2) cross-modal conditional probability. To investigate the computational principles underlying these response profiles 8 different models capturing various learning strategies were fit to the single-trial EEG data and compared via family-wise Bayesian model selection. A summary of the modelling results is depicted in figure 6.

**Figure 6.**
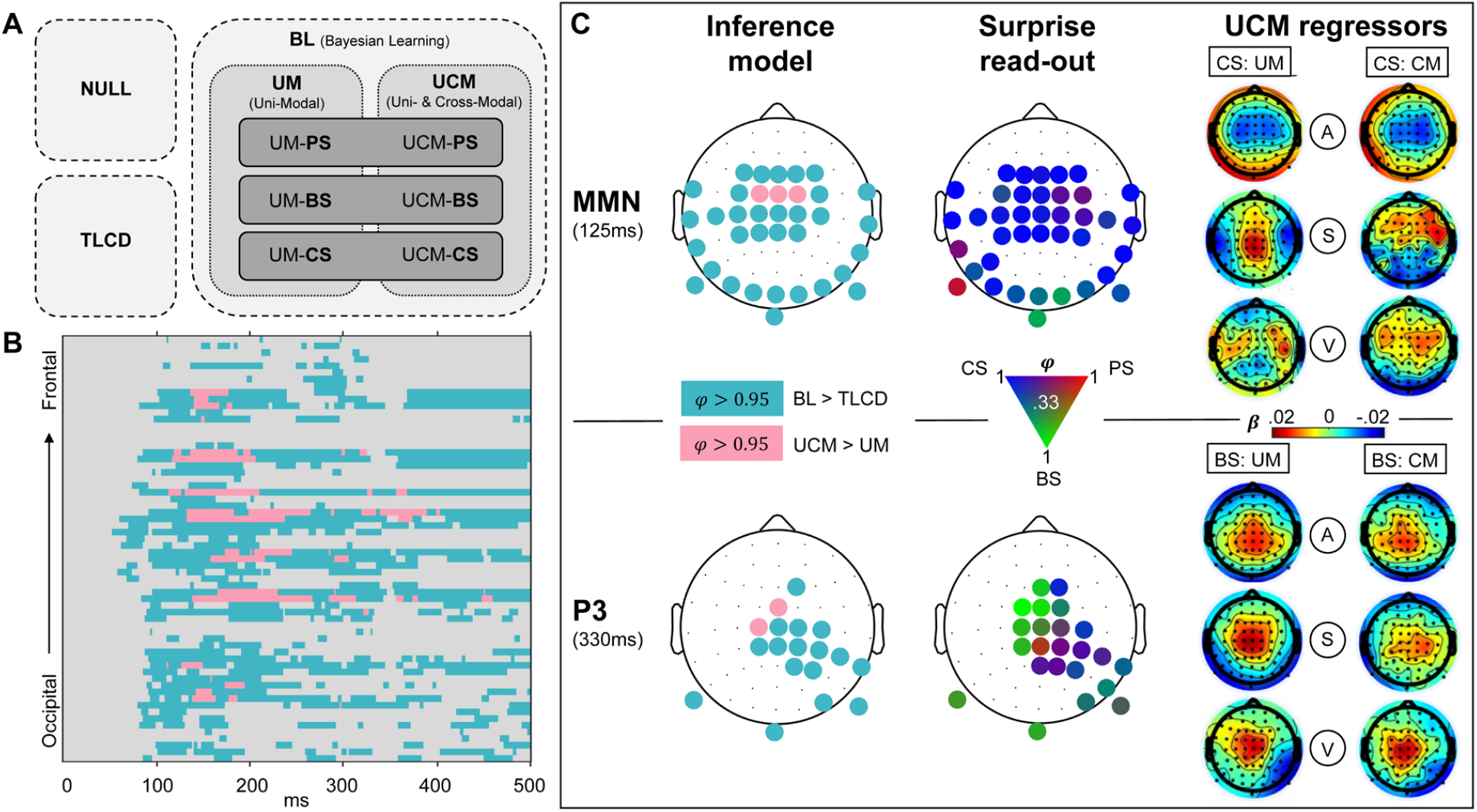
Modelling results. A) Schematic overview of models. Model comparison 1 (light-gray box, dashed contour): Null model family (NULL), train length dependent change detection model family (TLCD) and Bayesian learning model family (BL). Comparison 2 (gray box, dotted contour): Uni-modal regression model family (UM), cross-modal regression model family (UCM). Comparison 3 (dark-gray box, line contour): Read-out model family comparison of predictive surprise family (PS), Bayesian surprise family (BS) and confidence-corrected surprise family (CS). B) Results of comparison 1 and 2 shown for all electrodes and post-stimulus time points. Color depicts exceedance probability (EP) *φ* > 0.95. Light-blue=BL>TLCD, pink=UCM>UM. C) Topography of modeling results at time windows of MMN (top row) and P3 (bottom row). Left column: Results of comparison 1 (same colors as (B), depicting *φ* > 0.95). Middle column: Results of comparison 3. EPs between 0.33 and 1 of the three surprise functions are represented by a continuous 3-dimensional RGB scale (red=predictive surprise (PS); green=Bayesian surprise (BS); blue=confidence-corrected surprise (CS)). Right column: Beta estimates of the model regressors of the UCM model (regressors: A=auditory; S=somatosensory; V=visual; CM=cross-modal; UM=uni-modal) for CS read-out models (top) and BS read-out models (bottom).

The first model comparison aimed to further investigate observation (1) and the question whether the observed parametric modulation of standard and deviant EEG responses merely reflects a combination of neuronal adaptation and change detection dynamics or if the observed response patterns are indicative of an underlying generative model engaged in probabilistic inference. To this end, we ran family wise Bayesian model selection which is schematically depicted in figure 6A. The first comparison concerned the train length dependent change detection model (TLCD), a model family containing all Bayesian learning models, and a null model. In the fronto-central, temporal, occipital and central electrodes showing MMN and P3 effects, the model comparison shows strong evidence in favor of the BL model family with an exceedance probability *φ* > 0.95 from ∼70ms onward. On the other hand, the TLCD model did not exceed *φ* > 0.95 for any electrode or time point. Therefore, the TLCD model was disregarded at this point, and we focussed further investigation on the different BL models.

The second comparison set out to investigate observation (2) and evaluate the contribution of stimulus alternation tracking conditional on multi-modal configurations beyond uni-modal transition probability inference. Within the electrodes and time-points with sufficient evidence for Bayesian learning signatures (as established by the first model comparison), a comparison of a purely uni-modal model family with a cross-modally informed model family (UCM) was performed. Those electrodes and time-points where the additional inclusion of cross-modal regressors (UCM models) provided better model fits than purely uni-modal models are highlighted in figure 6B and C. The UCM family outperforms the UM family at central and fronto-central electrodes at ∼100-400ms (with *φ* > 0.95).

Inspection of the beta estimates of auditory, somatosensory and visual regressors of the UCM regression models shows that the beta maps of the uni-modal predictors of the model resemble the ERP mismatch topographies of the respective modalities (depicted in figure 6C). The cross-modal predictor, on the other hand, rather shows (fronto-)central activations which appear to resemble frontal aspects of the respective auditory, somatosensory and visual MMRs.

The third comparison concerned the three surprise measures used as read-out functions for the probabilistic models. Overall, the family comparison does not show overwhelming evidence for any specific surprise function as only few electrodes reach exceedance probabilities of *φ* > 0.95. Nevertheless, a tendency of the MMN and the P3 to reflect different surprise dynamics can be observed. Although around the time of the MMN, only some electrodes show *φ* > 0.95 in favour of confidence-corrected surprise, inspection of the topographies without *φ* thresholding (as depicted in figure 6C) shows CS to be dominant throughout the spatio-temporal range of the MMN (as suggested by higher EPs compared to BS and PS). On the other hand, at the time of the P3, Bayesian surprise appears to be the dominant surprise computation with multiple (fronto-) central electrodes showing *φ* > 0.95. Overall, the surprise comparison provides some evidence for a reflection of CS dynamics in the earlier mismatch signals around the time of the MMNs and suggests a tendency of the P3 to reflect BS dynamics.

In a final analysis, the optimal observation integration parameter *τ* was inspected. For each modality, the significant MMN clusters of the ERP analyses were used to inspect the optimal integration window of the regression models. For the UM regression model, highly similar optimal integration parameters were found within the electrodes and time-points of the different MMN clusters with no significant difference between the modalities. The optimal integration parameters were found to correspond to windows of stimulus integration with a half-life of (50% weighting at) around 5 to 20 stimuli. The same range of stimulus integration was found for the UCM regression model. Overall, confidence-corrected surprise models tended to have higher integration windows (∼10-20 stimuli) compared to Bayesian surprise models (∼5-10 stimuli).

## Discussion

The present study set out to compare mismatch signals in response to tri-modal sequence processing in the auditory, somatosensory and visual modalities and to investigate influences of predictive cross-modal information. We found comparable but modality specific signatures of MMN-like early mismatch processing between 100-200ms in all three modalities, which were source localized to their respective sensory specific cortices and shared right lateralized frontal sources. An additional cross-modal signature of mismatch processing was found in the P3 MMR for which a common network with frontal dominance was identified. With exception of the visual MMN, both mismatch signals (MMN and P3) show parametric modulation by stimulus train length driven by reciprocal tendencies of standards and deviants across modalities. Strikingly, standard and deviant responses within the cross-modal P3 cluster were sensitive to predictive information carried by the tri-modal stimulus configuration. Comparisons of computational models indicated that Bayesian learning models, tracking transitions between observations, captured the observed dynamics of single-trial responses to the roving stimulus sequences better than a static model reflecting train length dependent change detection. Moreover, a BL model which additionally captured cross-modal conditional dependence of stimulus alternation outperformed a purely uni-modal BL model primarily at central electrodes. The comparison of different read-out functions for the BL models provides tentative evidence that the early MMN may reflect dynamics of confidence-corrected surprise whereas later P3 MMRs seem to reflect dynamics of Bayesian surprise.

### Modality specific mismatch signatures in response to tri-modal roving stimuli

By using a novel tri-modal roving stimulus sequence originating from an underlying Markov process of state transitions, we were able to elicit and extract unique EEG signatures in each of the three sensory modalities (auditory, somatosensory and visual).

Of the EEG mismatch signatures, the auditory MMN is one of the most widely researched responses to deviation from an established stimulus regularity (Näätänen 1978; Winkler 2009). Contrasting responses to standard and deviant stimuli of the auditory sequence in the current study resulted in the expected fronto-central MMN signature with more negative responses to deviants compared to standards. The extent of the MMN might suggest an underlying negative mismatch component as proposed by Näätänen (2005)), which drives a more negative going ERP around the N1, extending beyond the P2 component. Such post-N1 effects of the MMN have been suggested as markers of a “genuine” mismatch component in contrast to confounds by stimulus properties modulating auditory ERP components (Näätänen 2007) and might speak against pure N1 adaptation (as suggested by May 2004; Jääskiläinen 2004; May 1999).

The somatosensory equivalent to the auditory MMN (sMMN) reported in the current study shows negative polarity at bilateral temporal electrodes and corresponding central positivity. The sMMN likely reflects an enhanced N140 component, as suggested by Kekoni et al. (1997). However, most previous sMMN studies used oddball paradigms where some critical discussion revolves around the distinction of the sMMN from an N140 modulation by stimulus properties alone. Here, we report an sMMN around the N140 which can be assumed to be independent of stimulus confounds due to the reversed roles of standard and deviant stimuli in the roving paradigm. Although several previous studies have reported somatosensory mismatch responses, conflicting evidence exists regarding the exact components that may constitute an equivalent to the auditory MMN. Some studies report a more fronto-centrally oriented negativity (Kekoni 1997; Spackman 2007; 2010; Shen 2018) or observed such pronounced central positivity that they were led to conclude that it is in fact the central positivity that should be considered the somatosensory equivalent of the aMMN (Shinozaki 1998; Akatsuka 2005). However, some evidence appears to converge on a temporally centred negativity with corresponding central positivity as the primary sMMN around 140ms (Ostwald 2012; Gijsen 2021).

While the auditory and somatosensory MMN’s in the current study were found to be highly comparable in their signal strength, their hypothesized counterpart in the visual modality showed a comparatively weaker response. Nevertheless, we found a significant visual MMN (vMMN) at occipital electrodes extending to temporal electrodes within a time window of 100-200ms post stimulus, with corresponding (fronto-)central positivity. This observation is in line with previous research reporting posterior (Urakawa 2010; Kimura 2010; Clery 2013) and temporal (Hesenfeld 2003; Kuldkepp 2013) patterns of vMMN with corresponding central positivity (Czigler 2006; Cleary 2013; File 2017).

### Neuronal generators of MMN signatures

Source reconstruction analyses were used to identify underlying neuronal generators of the modality specific MMN signatures. Interestingly, for each sensory modality, we found generators in the primary and higher order sensory cortices as well as additional frontal generators in inferior frontal gyrus (IFG) and middle frontal gyrus (MFG).

The sensory specific neuronal sources underlying the auditory MMN were identified as bilateral auditory cortex with a dominance in hierarchically higher auditory areas. With an additional modality independent contribution of right lateralized frontal sources, this set of neuronal generators identified for the aMMN is in line with previous research suggesting primary auditory cortex and higher auditory areas in superior temporal sulcus (STG) as well as right IFG as underlying the aMMN (Opitz 2002; Molholm 2005; Näätänen 2005; Garrido 2008; Garrido 2009a) with consideration of an additional frontal generator in MFG (Deouell 2007).

The sources underlying the sMMN were identified in the current study as primary (S1) and secondary (S2) somatosensory cortices with additional frontal generators in right IFG and MFG. This finding is in accordance with previous research showing a combined response of S1 and S2 to underlie the sMMN (Akatsuka 2007a; 2007b; Spackman 2010; Butler 2012; Ostwald 2012; Naeije 2016; 2018; Andersen 2019; Gijsen 2021) in combination with involvement of (inferior) frontal regions (Huang 2005; Ostwald 2012; Allen 2016; Fardo 2017; Downar 2000).

For the visual modality, we identified sources in visual areas (V1-V4) and additional frontal activations in IFG and MFG as the neuronal generators underlying the vMMN. Previous studies have shown similar combinations of visual and prefrontal areas (Yucel 2007; Kimura 2010; 2011; 2012; Urakawa 2010) and have particularly highlighted the IFG as a frontal generator of the vMMN (Downar 2000; Hedge 2015). Similarly, an fMRI study of perceptual sequence learning in the visual system has shown right lateralized prefrontal activation in addition to activations in visual cortex in response to regularity violations (Huettel 2002). Yet another study has suggested a role for right prefrontal areas in interaction with hierarchically lower visual areas for the prediction of visual events (Kimura 2012), all in line with our results.

Overall, our finding of inferior and middle frontal sources for the MMN in all three modalities provides further evidence for a modality independent role for these generators as previously suggested by Downar et al. (2000). As such, these modality-independent frontal generators might reflect higher stages of a predictive hierarchy working across modalities in interaction with lower modality specific regions, as previously suggested primarily for the auditory modality (Garrido 2009b).

### Modulation of the MMN by stimulus repetition

An important feature of the MMN which theories of its generation have aimed to account for is its sensitivity to stimulus repetition. The MMN is known to increase with prior repetition of standards (Sams 1983; Näätänen 1992; Imada 1993; Javitt 1998). Correspondingly, in the current study we find a significant increase of auditory and somatosensory MMN with the length of the preceding stimulus train as well as a comparable tendency for the visual MMN. Moreover, we show that this increase was driven by a reciprocal negative modulation of deviant and positive modulation of standard responses, suggesting a combined influence of repetition dependent change detection and dynamics akin to stimulus adaptation.

The observed positive modulation of standard responses, particularly in the auditory modality, is in line with the repetition positivity account of Baldeweg and colleagues (2004; 2006; 2007; Haenschel 2005). In the auditory modality, repetition positivity has been isolated as a positive slow wave that accounts for repetition-dependent increases of auditory ERPs up to the P2 component (Haenschel 2005). With regards to its functional role, it has been argued to reflect auditory sensory memory trace formation (Baldeweg 2004; Costa-Faidella 2011a; Costa-Faidella 2011b). Interestingly, MMN studies using the oddball paradigm often report an increasing MMN with standard repetition without further dissecting the contributions from standard and deviant dynamics. A contribution of the standard repetition positivity appears to be particularly dominant in roving stimulus paradigms (Cooper 2013), potentially because a memory trace of the standard stimulus identity must be re-established after each change of roles for standard and deviant stimuli. It has even been suggested that the memory trace dynamics of the standard observed in response to roving oddball sequences might in fact be the primary driver of train length effects on MMN amplitudes (Baldeweg 2004; Haenschel 2005; Costa-Faidella 2011a; Costa-Faidella 2011b). Importantly, although some evidence exists to suggest an additional role for train length dependent deviant modulation also in roving paradigms (Cowan 1993; Haenschel 2005), a dissection of combined standard and deviant contributions as performed here is rarely described.

Similar to the aMMN, we found the sMMN to be modulated by stimulus repetition. An early repetition positivity effect in the responses to standards was observed prior to 100ms indicating comparable sensory adaptation dynamics as described for the aMMN. Subsequently, the negative deviant and sMMN responses increase with repetition around the N140 (i.e., around the sMMN peak). While somatosensory deviant responses have previously been shown to decrease with increasing stimulus probability (Akatsuka 2007), only few other studies have reported sensitivity of the sMMN to stimulus repetition. Interestingly, in our previous study on somatosensory MMRs (Gijsen 2021) we report the same reciprocal pattern found here: Negative modulation of the deviant and positive modulation of the standard response which result in an increase of the sMMN amplitude with stimulus train length. In the visual modality, a comparable train length effect to auditory and somatosensory modalities was observed but did not reach statistical significance in the vMMN time window. Given the overall weaker response in the current study for vMMN this might not be surprising. Moreover, discussions about the repetition modulation of vMMN responses are often based on findings concerning the auditory system rather than direct findings in the visual modality. While sensory adaptation to stimulus repetition is generally found throughout the visual system (e.g. Grill-Spector 2006; Clifford 2007) it is rarely directly reported in visual MMN studies (but see Kremláček 2016). Overall, the visual MMN literature seems to suggest that the vMMN may be a rather unstable phenomenon. In fact, by controlling for confounding effects, one study has called the existence of the vMMN for low level features such as the ones used here into question entirely (Male 2020). The vMMN appears to show a much less pronounced spatiotemporal pattern than auditory and somatosensory equivalents, which is reflected in larger variance in the reported topographies and time windows in studies investigating vMMN (but see *Limitations* for a discussion of alternative explanations regarding the current study).

### MMN as a signature of predictive processing

Recent research supports the view that Bayesian perceptual learning mechanisms underlie the generation of mismatch responses such as the MMN (Friston 2005; 2010; Garrido 2009b). Given the proposal of Bayesian inference and predictive processing as universal principles of perception and perceptual learning in the brain (Friston 2005; 2010), comparable mismatch responses are expected to be found across sensory modalities. Evidence for the predictive nature of mismatch responses, akin to key findings from the auditory modality, is for instance given by studies showing somatosensory (Naeije 2018; Andersen 2019) and visual (Czigler 2006; Kok 2014) MMN in response to predicted but omitted stimuli. Moreover, Ostwald et al. (2012) and Gijsen et al. (2021) have shown that single trial somatosensory MMN and P3 MMRs can be accounted for in terms of surprise signatures of Bayesian inference models tracking stimulus transitions. Similarly, the vMMN has been described as a signature of predictive processing (Kimura 2011; Stefanics 2014), signaling prediction error instead of basic change detection (Stefanics 2018).

Correspondingly, we found comparable mismatch signatures in auditory, somatosensory and visual modalities. The train length effects observed in our study across modalities have previously been related to predictive processing. Repetition positivity in the auditory modality has been interpreted as a reflection of repetition suppression, resulting from fulfilled prediction (Aukstulewicz 2016; Baldeweg 2006, 2007; Costa-Faidella 2011a; Costa-Faidella 2011b). A corresponding negative modulation of deviant responses on the other hand, would signal a failure to suppress prediction error after violation of the regularity established by the current stimulus train. Under such a view, longer trains of repetitions lead to higher precision in the probability estimate which in turn results in a scaling of the prediction error in response to prediction violation (Friston 2005; 2009; Aukstulewicz 2016). In line with these hypotheses, Garrido and colleagues (2008; 2009a) used dynamic causal modelling (DCM) to show that the MMN elicited in a roving stimulus paradigm is best explained by the combined dynamics of auditory adaptation and model adjustment. Their network, proposed to underlie MMN generation, was set up as an implementation of hierarchical predictive processing involving bottom-up signals from auditory cortex and top-down modulations by inferior frontal cortex. Similarly, another DCM study proposed a predictive coding model of pain processing in response to somatosensory oddball sequences, highlighting the role of inferior frontal cortex in top-down modulations of somatosensory potentials (Fardo 2017). As we find involvement of such modality specific sensory and modality independent frontal areas for MMN responses across modalities, our results suggest comparable roles for these sources in a predictive hierarchy.

### P3 mismatch responses reflect cross-modal processing

In addition to the modality specific MMN responses, deviants in all three modalities elicited a late positive mismatch component in the P3 time window. Despite differences in the exact latency and extent of this response between modalities, we identified a common mismatch cluster from 300-350ms in central electrodes, followed by a slightly more posterior cluster extending from 400-600ms. Particularly the earlier cluster may correspond to the well-known P3a response, which peaks at around 300ms after change-onset at (fronto-)central electrodes and is thought to be elicited regardless of sensory modality (Escera 2000; Friedman 2001; Knight 1998; Schroeger 1996; Polich 2007).

The P3a is closely related to the MMN as they are both elicited during active and passive perception of repeated stimuli interrupted by infrequent stimulus deviations (Polich 2007; Schroeger 2015). While the P3a has been initially related to attentional switches to task-irrelevant but salient stimulus features (Escera 2000; Friedman 2001; Polich 2007), more recent accounts suggest that the MMN and P3a might reflect two stages of a predictive hierarchy, each representing (potentially differentiable) prediction error responses (Wacongne 2011; Schroeger 2015). Similar to the MMN, P3 responses are known to be modulated by stimulus probability (Duncan-Johnson 1977) and can be elicited by unexpected stimulus repetitions (Squires 1976; Duncan 2009) and omissions of predicted sound stimuli (Sutton 1967; Prete 2022), which provides compelling evidence for a role of the P3 in predictive processing. Similar to the MMN responses described above, we found the individual P3 MMR responses in all three modalities to show reciprocal modulations of standards and deviants by stimulus repetition, which has previously only been reported for the auditory modality (Bendixen 2007). This sensitivity to stimulus repetition of mismatch responses in early and late time-windows has been interpreted in terms of regularity and rule extraction in the auditory modality (Bendixen 2007) and is in line with an account of repetition suppression over and above early sensory adaptation.

The MMN and P3 MMR have been shown to be differentially modulated by higher order predictability. The P3 is reduced by the presentation of visual cues preceding an auditory deviant, while the MMN is not affected by the same top-down predictability (Ritter 1999; Sussman 2003; Horvath 2012). Similarly, explicit top-down knowledge of sequence regularities has been shown to reduce the P3, while leaving the MMN unaffected (Max 2015). It has thus been suggested that the P3 reflects a higher-level deviance detection system concerned with the significance of the stimulus in providing new information for the system (Horvath 2008). Interestingly, a recent study investigating mismatch responses to different auditory features showed that while the MMN response in an earlier (classical) time window was generally affected by regularity violations, only the later response (P3 range) contained information about the specific features that were violated (An 2021). Furthermore, computational studies indicate that P3 responses reflect specific quantities of unexpectedness as well as updates to a prior belief (Jepma 2017; Kolossa 2015).

Overall, current research provides evidence for the view that the MMN reflects prediction errors at earlier hierarchical stages, primarily concerned with more local regularity extraction, whereas P3 responses reflect more global rule violations which require a certain level of abstraction and information integration (Wacongne 2011; Bekinschtein 2009; Winkler 2005). Our findings of a sensitivity of the P3 response to cross-modal predictive information carried by the multi-modal configuration of the stimulus sequence further supports such a view. Across modalities, we found an increased P3 response to mispredicted compared to predicted or unpredictable stimuli, regardless of their role as standards or deviants. Generally, the P3 deviant response in the current study likely reflects a (unsigned) prediction error to a local regularity established by stimulus repetition. However, increased P3 responses to mispredicted stimuli indicate additional violations of global, cross-modal predictions which are extracted from multi-modal context information.

The observed pattern suggests influences of precision weighting on prediction errors (Friston 2009). In case of both predicted and mispredicted stimuli, the cross-modal predictive context allows for more precise predictions (i.e. high prior precision) than in case of the unpredictable stimuli (low prior precision). Under such an interpretation the precision for mispredicted deviants is high, resulting in a pronounced prediction error response. Since the precision for predicted deviants is also high, the resulting prediction error response is low because the stimulus was suppressed. Even though the size of prediction error to unpredictable deviants could generally be expected in between those of predicted and mispredicted deviants, the observed response is low (similar to that of a predicted deviant), because the prior precision in this context is low. This interpretation is in line with the fact that no significant difference was found between predicted and unpredictable deviants. A similar modulation of multi-modal predictability is found for the P3 response to standards. However, interestingly, in case of the standards, the response to predicted stimuli is significantly lower than to unpredictable stimuli. This difference between standards and deviants could be due to the fact that deviants are generally surprising, even if they are more predictable in terms of their cross-modal configuration. Standards, on the other hand, are generally predicted to occur (high precision) which might result in a pronounced suppression of prediction error in case they are additionally cross-modally predicted.

The interpretation of the common P3 cluster as a cross-modal P3a response sensitive to multi-modal predictive information is further supported by our source localization results, which particularly indicate prefrontal regions such as the medial frontal, inferior frontal and anterior cingulate cortex as sources of the P3 MMR. Although notoriously diverse, previous research on P3 sources has identified a fronto-parietal network of generators, particularly highlighting the role of prefrontal and anterior cingulate regions in generating the P3 novelty response (P3a) (Linden 2005; Polich 2007), whereas parietal regions are presumed to be more involved in task-related P3b responses. The identified sources have been shown to be involved in a fronto-parietal network relevant for the supra-modal processing of stimulus transitions and deviance detection (Downar 2000; Huang 2005). Similarly, a fronto-parietal attention network (Corbetta 2002) has been shown to be involved in oddball processing in the auditory and visual modalities (Kim 2014). The network consists of two functionally and anatomically distinct parts which closely interact (Vossel 2014). While the dorsal part of the network is believed to be involved in the allocation of top-down, endogenous attention (e.g. triggered by predictive information), the ventral part is involved in bottom-up, exogenous attention allocation and thus, processing of unexpected stimuli. Importantly, it has been shown that this network operates supra-modally to facilitate processing of information from multi-modal events (Macaluso 2005; 2010). Thus, the predictive information in the multi-modal sequences presented in the current study may be processed in such a fronto-parietal network to aid the perception of multi-modal stimulus streams. Future research would benefit from studies further investigating such multi-modal probabilistic sequences with higher spatial resolution to inform these proposed interpretations.

### Modelling single-trial EEG responses as signatures of Bayesian inference

Given the results of the average- and GLM-based EEG analyses, we aimed to test if the observed modulations of standards, deviants and MMRs by local (train length) and global (cross-modal predictability) sequence properties could be captured by signatures of Bayesian inference. To this end, we compared a simple train length dependent change detection (TLCD) model to families of Bayesian learning (BL) models capturing different aspects of the sequence statistics. In light of the literature discussed above we hypothesized that BL models would outperform the TLCD model in explaining the recorded mismatch responses.

Overall, the BL models outperformed the static TLCD model in all electrodes in the MMN and P3 clusters indicating that these responses reflect dynamics beyond the basic repetition effects observed in the ERP analyses. This result provides evidence to suggest that the MMN and P3 MMR capture the trial-to-trial dynamics of Bayesian inference and are thus markers of probabilistic sequence processing in the brain.

Within the family of BL models, we found that a cross-modally informed model (UCM), tracking cross-modal conditional dependencies between modalities in addition to uni-modal transitions, outperformed a purely uni-modal transition probability model (UM) at central electrodes within an early and a late time-window. The cross-modal effects in the late time-window are directly in line with the sensitivity of the P3 cluster to cross-modal predictability discussed above and support an interpretation of P3 mismatch responses to reflect signatures of cross-modal Bayesian inference. Given that cross-modal learning was not explicitly instructed or task-relevant, the results are compatible with the view that the brain is sensitive to cross-modal information by default (Ghazanfanar 2006; Driver 2008) and that processing multi-modal information might be appropriately captured by Bayesian inference (Kording 2007; Shams 2022). Interestingly, however, an earlier cross-modal effect was found prior to 300ms which was not reflected in the GLM results, suggesting that potential modulations of MMN signatures by predictability manifest in the dynamics of single trial surprise signals but not in significant mean differences between predictability conditions. Since the earlier cross-modal effect observed in the modelling results was primarily confined to central and fronto-central electrodes it may be related to activity of the frontal generators of the MMN. As discussed above, the frontal cortex is assumed to be involved in MMN generation (Deouell 2007) in interaction with hierarchically lower sensory sources and has been hypothesized to form top-down predictions about incoming sensory stimuli (Garrido 2008; 2009a, 2009b). This assumption is further supported by our source reconstruction results which show modality independent frontal generators in addition to sensory specific regions to underlie the MMN in auditory, somatosensory and visual modalities.

Regarding the surprise read-out functions of the BL models, we find a slight dominance of confidence-corrected surprise (CS) in earlier mismatch signatures prior to 200ms, while the late clusters tend to reflect Bayesian surprise (BS). This is well in line with our previous study performed in the somatosensory modality (Gijsen 2021) and other studies have similarly reported a reflection of BS in P3 mismatch responses (Ostwald 2012; Kolossa 2015; Mars 2008; Seer 2016). Given their differences in reading out the probability estimates of the Bayesian observer, the different surprise signatures in the MMN and P3 MMR might provide some insight into their respective computational roles. CS has been categorized as an instantiation of puzzlement surprise (Faraji 2018) reflecting a mismatch between sensory input and internal model belief which is additionally scaled by belief commitment. Low-probability events are thus more surprising if commitment to the belief (of this estimate) is high. BS reflects incorporation of new information, quantifying an update to the generative model and has been categorized as enlightenment surprise (Faraji 2018). Accordingly, the MMN may be considered a marker of prediction error scaled by belief commitment, whereas the P3 may reflect the subsequent update of the predictive model.

### Limitations

Although we gained valuable insights into the commonalities and differences between mismatch responses in different modalities, our study faces certain limitations in its implementation and scope. First, although reports of weak vMMN responses can be found in the literature, an alternative explanation may lie in the stimulation protocol used in the current study. Our visual stimuli consisted of bilateral flash stimuli with two different intensities, which were presented in the periphery of the visual field. Since, to our knowledge, no other study has used visual flash stimuli to elicit vMMN, our results are not directly comparable to previous research. Moreover, due to the retinotopic organisation of the visual cortex (Sereno 1995; Horton 1991), a “far peripheral” placement (i.e. >60 degrees; Strassburger 2011) of the LED’s results in the activation of (primary) visual areas folded deep inside the cortex, in the calcarine sulcus between the hemispheres. It is therefore possible that the visual mismatch responses were not weaker per se but were merely harder to detect by means of EEG.

Further, our results concerning the comparison of the surprise read-out functions provide some indications of the computational roles for early and late MMRs, which are in line with previous research. However, it should be noted that the current study was not specifically designed to investigate their (nuanced) differences. The inclusion of three read-out functions primarily served the purpose of avoiding bias in the comparison of the Bayesian learning models by prior choice of the read-out. To this end, the most prominent surprise read-out functions used in the literature were included. Research would benefit from future studies specifically designed to compare different surprise measures without the manipulation of other aspects of the underlying models. A valuable overview and suggested experiments for that purpose have been recently provided by Modirshanechi et al. (2022).

## Conclusion

With the current study we provide evidence for modality specific and modality independent aspects of mismatch responses in audition, somatosensation and vision resulting from a simultaneous stream of tri-modal roving stimulus sequences. Our results suggest that responses to stimulus transitions in all three modalities are based on an interaction of hierarchically lower, modality specific areas with hierarchically higher, modality independent frontal areas. We show that similar dynamics underlie these mismatch responses which likely reflect predictive processing and Bayesian inference on uni-modal and multi-modal sensory input streams.

## Acknowledgement

The authors would like to thank the HPC Service of ZEDAT, Freie Universität Berlin, for computing time.

